# Genesis and Gappa: Processing, Analyzing and Visualizing Phylogenetic (Placement) Data

**DOI:** 10.1101/647958

**Authors:** Lucas Czech, Pierre Barbera, Alexandros Stamatakis

**Affiliations:** Computational Molecular Evolution Group, Heidelberg Institute for Theoretical Studies, Heidelberg, Germany; Institute for Theoretical Informatics, Karlsruhe Institute of Technology, Karlsruhe, Germany

**Keywords:** Metagenomic Analysis Tool, Data Visualization, Phylogenetic Trees, Phylogenetic Placement, Evolutionary Placement, Software, Library

## Abstract

We present GENESIS, a library for working with phylogenetic data, and GAPPA, an accompanying command line tool for conducting typical analyses on such data. The tools target phylogenetic trees and phylogenetic placements, sequences, taxonomies, and other relevant data types, offer high-level simplicity as well as low-level customizability, and are computationally efficient, well-tested, and field-proven.

**Availability and Implementation:** Both GENESIS and GAPPA are written in modern C++11, and are freely available under GPLv3 at http://github.com/lczech/genesis and http://github.com/lczech/gappa.

## 1. Introduction

The necessity of computation in biology, and in (metagenomic) sequence analysis in particular, has long been acknowledged (1). In phylogenetics, for example, there is a plethora of scientific software for analyzing data, covering tasks such as sequence alignment (2), phylogenetic tree inference (3), and diverse types of downstream analyses (4). Furthermore, in metagenomics, a key task is the taxonomic identification of sequences obtained from microbial environments. An increasingly popular method for this is phylogenetic (or evolutionary) placement, which can classify large numbers of (meta-)genomic sequences with respect to a given reference phylogeny. Common tools for phylogenetic placement are pplacer (5), RAxML-EPA (6), as well as the more recent and more scalable EPA-ng (7). The result of a phylogenetic placement can be understood as a distribution of sequences over the reference tree, which allows to examine the composition of microbial communities, and to derive biological and ecological insights (8, 9).

Here, we introduce genesis, a library for working with phylogenetic data, as well as gappa, a command line tool for typical analyses of such data. They focus on phylogenetic trees and phylogenetic placements, but also offer various additional functionality. Combined, they allow to analyze as well as visualize phylogenetic (placement) data with existing methods and to experiment with and develop novel ideas.

To maximize usability of our tools, our implementation is guided by the following design objectives: (a) Most users require a fast and simple application for analysing their data, (b) some power users desire customization, e. g., via scripting, (c) developers require a flexible toolkit for rapid prototyping, and (d) with the on-going data growth, the implementation needs to be scalable and efficient with respect to memory and execution times. To this end, genesis and gappa are written in `C++11`, relying on a modern, modular, and function-centric software design.

We evaluate the code quality, the runtime behavior, and the memory requirements for conducting typical tasks such as file parsing and data processing in the Supplementary Text. An exemplary benchmark for reading `newick` files is shown in Fig. 1. We find that genesis has the overall best code quality score (10) compared to other scientific codes written in `C` or `C++`. It is also consistently faster than all evaluated `Python` and `R` libraries in our tests. Furthermore, gappa is faster and more memory efficient than its main competitor guppy in almost all tests and, most importantly, it scales better on larger datasets in all benchmarks.

**Fig. 1.**
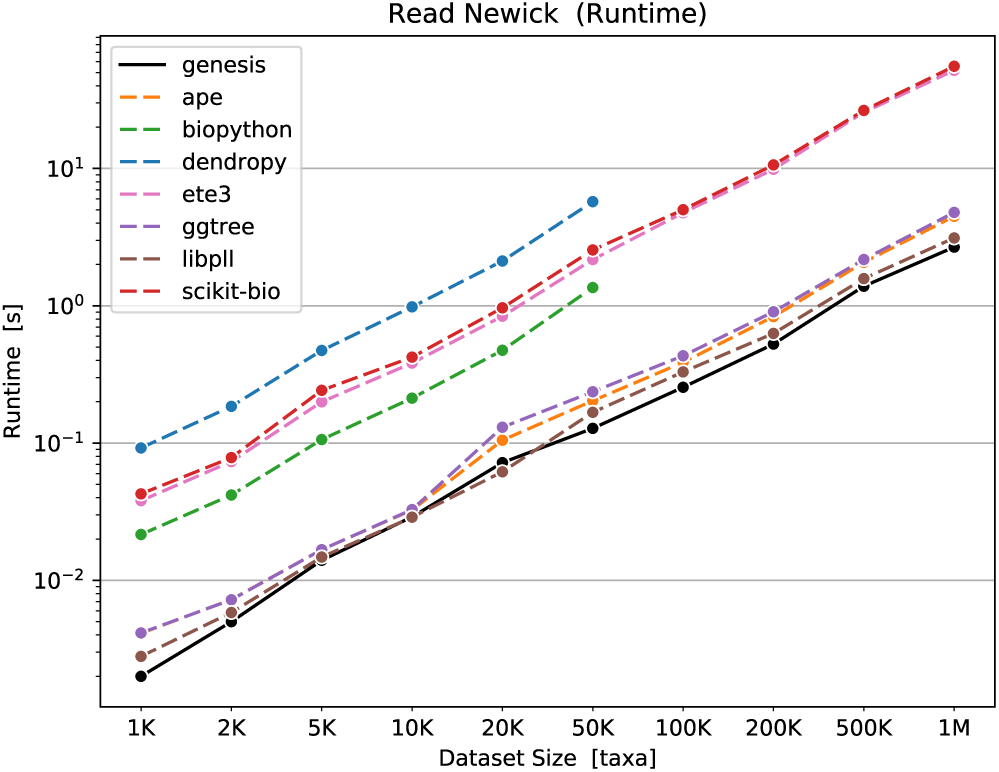
Runtimes for reading `newick` files with 1K to 1M taxa (tip/leaf nodes) and a randomly generated topology, using a variety of diffent libraries and tools. See the Supplementary Text for details.

## 2. Features of Genesis

genesis is a highly flexible library for reading, manipulating, and evaluating phylogenetic data with a simple and straight-forward API. Typical tasks such as parsing and writing files, iterating over the elements of a data structure, and other frequently used functions are mostly one-liners that integrate well with modern `C++`. The library is multi-threaded, allowing for fully leveraging multi-core systems for scalable processing of large datasets. The functionality is divided into loosely coupled modules, which are organized in `C++` namespaces. We briefly describe them in the following.

### Phylogenetic Trees

Phylogenetic trees are implemented via a pointer-based data structure that enables fast and flexible operations, and allows to store arbitrary data at the nodes *and* at the edges. The trees may contain multifurcations and may have a designated root node. Trees can be parsed from `newick` files and be written to `newick`, `phyloxml`, and `nexus` files, again including support for arbitrary edge and node annotations. Traversing the tree starting from an arbitrary node in, e. g., post-order, pre-order, or level-order can be accomplished via simple `for` loops:

~~~
// Read a tree from a Newick file.
Tree tree = Common Tree Newick Reader (). read (
  from_ file (“ path / to/ tree. newick “)
);
// Traverse tree in preorder, print node names.
for (auto it : preorder (tree)) {
  auto & data = it. node (). data < Common NodeData >();
  std :: cout << data. name << “\ n”;
}
~~~

Functionality for manipulating trees, finding lowest common ancestors of nodes (e. g., using the Euler tour technique of (11)) or paths between nodes, calculating distances between nodes, testing monophyly of clades, obtaining a bitset representation of the bipartitions/splits of the tree, and many other standard tasks is provided. Furthermore, functions for drawing rectangular and circular phylograms or cladograms to `svg` files, using individual custom edge colors and node shapes, are provided for creating publication quality figures.

### Phylogenetic Placements

Handling phylogenetic placement data constitutes a primary focus of genesis. Placement data is usually stored in so-called `jplace` files (12). Our implementation offers low-level functions for reading, writing, filtering, merging, and otherwise manipulating these data, as well as high-level functions for distance calculations (13), Edge PCA and Squash Clustering (14), and Phylogenetic *k*-means clustering (9), among others. Advanced functions for analysing and visualizing the data are implemented as well, for instance, our adaptation of Phylofactorization to phylogenetic placement data (9, 15). Lastly, we offer a simple simulator for generating random placement data (e.g., for testing).

To the best of our knowledge, competing software that can parse placement data in form of `jplace` files (BoSSA (16), ggtree (17), or iTOL (18)) merely offers some very basic analyses and visualizations, such as displaying the distribution of placed sequences on the reference phylogeny, but does not offer the wide functionality range of genesis.

### Other Features

Sequences and alignments can be efficiently read from and written to `fasta` and `phylip` files; high-level functions for managing sequences include several methods for calculating consensus sequences, the entropy of sequence sets, and sequence re-labelling via hashes. Taxonomies and taxonomic paths (for example, “`Eukaryota;Alveolata; Apicomplexa`”) can be parsed from databases such as Silva (19, 20) or NCBI (21, 22) and stored in a hierarchical taxonomic data structure, again with the ability to store arbitrary meta-data at each taxon, and to traverse the taxonomy.

Furthermore, genesis supports several standard file formats such as `json`, `csv`, and `svg`. All input methods automatically and transparently handle `gzip`-compressed files. Moreover, a multitude of auxiliary functions and classes is provided: Matrices and dataframes to store data, statistical functions and histogram generation to examine such data, regression via the Generalized Linear Model (GLM), Multi-Dimensional Scaling (MDS), *k*-means clustering, an efficient bitvector implementation (e. g., used for the bitset representation of phylogenetic trees mentioned above), color support for handling gradients in plots, etc. The full list of functionality is available via the online documentation.

Lastly, genesis offers a simple architecture for scripting-like development, intended for rapid prototyping or small custom programs (e. g., convert some files or examine some data for a particular experiment).

## 3. Features of Gappa

The flexibility of a library such as genesis is primarily useful for method developers. For most users, it is however more convenient to offer a simple interface for typical, frequent tasks. To this end, we have developed the command line program gappa.

gappa implements and makes available the methods we presented in (8) and (9), such as: automatically obtaining a set of reference sequences from large databases, which can be used to infer a reference tree for phylogenetic placement; visualization tools to display the distribution of placements on the tree or to visualize per-branch correlations with meta-data features of environmental samples; analysis methods such as Phylogenetic *k*-means and Placement-Factorization for environmental samples.

gappa also contains re-implementations of a few prominent methods of guppy (5), as well as commands for sanitizing, filtering, and manipulating files in formats such as `jplace`, `newick`, or `fasta`, and a command for conducting a taxonomic assignment of phylogenetic placements (23).

As gappa internally relies on genesis, it is also efficient and scalable. Hence, gappa can also be considered as a collection of demo programs for using genesis, which might be helpful as a starting point for developers who intend to use our library. In comparison to guppy, we have observed speedups of several orders of magnitude and significantly lower memory requirements in general when processing large data volumes; see the Supplementary Text and (8) for details.

## 4. Conclusion

We presented genesis, a library for working with phylogenetic (placement) data and related data types, as well as gappa, a command line interface for analysis methods and common tasks related to phylogenetic placements. genesis and gappa already formed an integral part in several of our previous publications and programs (7–9, 24, 25), proving their flexibility and utility.

In future genesis releases, we intend to offer API bindings to `Python`, thus making the library more accessible to developers. In gappa, we will implement additional commands, in particular for working with phylogenetic placements, as well as re-implement the remaining commands of guppy, in order to facilitate analysis of larger data sets.

Both genesis and gappa are freely available under GPLv3 at http://github.com/lczech/genesis and http://github.com/lczech/gappa.

## Appendices

### Author’s Contributions

LC designed and implemented genesis. LC and PB implemented gappa. All authors helped to draft the manuscript. All authors read and approved the final manuscript.

## Acknowledgments

This work was financially supported by the Klaus Tschira Stiftung gGmbH Foundation in Heidelberg, Germany. We thank all members of the Exelixis Lab for their help and suggestions. The background music for this work was provided by Peter Gabriel and Phil Collins.

## Supplementary Text: Software Comparison

In this supplement, we compare genesis and gappa to competing software. First, we evaluate the runtime and memory requirements for some representative and frequent tasks, by comparing genesis to libraries and tools that have similar scope and functionality. Second, we compare gappa to the guppy tool by benchmarking some of their common functionality. guppy is part of the pplacer suite of programs [1], which were one of the first phylogenetic placement tools. Third, we evaluate the quality of the genesis code in comparison to other scientific software written in `C++` and `C`.

### 1 Runtime and Memory

We compare genesis and gappa to several competing libraries and tools. We focus on the most widely used alternative tools. The full list of competing software to which we compared genesis and gappa is provided in Table S1. Most general-purpose libraries for bioinformatics tasks, manipulation and (post-)analysis of genetic sequences and phylogenetic trees are written in interpreted languages such as `Python` or `R`. In order to also compare genesis to software written in a similar, compiled, language, we also included two other libraries developed by our lab, namely libpll [2] and the accompanying pll-modules [3]. They are written in `C`, and are the core libraries of the recent re-implementation of RAxML-ng [4]. We however note that these libraries are independed of genesis and do not share any code.

We compare several typical tasks for working with sequences, trees, and phylogenetic placements. As the specific scope of each library and tool differs, there is comparatively little overlap in functionality that can be used for benchmarking. Hence, we evaluated the runtime and memory requirements of genesis for the following general tasks:

- Parsing files in `fasta` format [5]; see Figure S1.
- Parsing files in `phylip` format [6]; see Figure S2.
- Calculating base frequencies on sets of sequences; see Figure S3.
- Parsing files in `newick` format [7]; see Figure S4.
- Calculating the pairwise patristic distance matrix on a given tree with branch lengths; see Figure S5.
- Parsing files in `jplace` format [8]; see Figure S6.

Furthermore, we evaluted the runtime and memory requirements of gappa for the following types of typical analyses of phylogenetic placement data:

- Calculate the EDPL of the queries in a placment sample [1]; see Figure S7.
- Calculate the Kantorovich-Rubinstein (KR) distance [9] between sets of placement samples; see Figure S8.
- Calculate the Edge Principal Components (Edge PCA) [10] of sets of placement samples; see Figure S9.
- Calculate the Squash Clustering [10] of sets of placement samples; see Figure S10.

We provide all scripts used for testing the tools and creating the plots shown here at https://github.com/lczech/genesis-gappa-paper. The tests were run on a 4-core laptop with 12 GB of main memory. We note that genesis and gappa offer parallel processing and computation. They can, for example read/parse multiple files simultaneously on distinct compute cores. We are not aware that any of the competing libraries and tools evaluated here transparently offer this feature. Hence, for a fair comparison, we also tested genesis and gappa on a single core only.

In summary, genesis outperforms *all* tools written in `Python` and `R`. In most cases it is also more memory-efficient. The runtime and memory requirements for libpll/pll-modules in comparison to genesis show that each of the tools performs better at specific tasks, depending on how much effort was invested into the optimization of the respective method. Based on a careful code inspection, we believe that the tasks were each software performs best are implemented close to optimal regarding runtime and/or memory.

Furthermore, gappa generally outperforms guppy in all tests, both in terms of runtime and memory usage. There is only one type of analysis (Edge PCA, Figure S9) where guppy has a slight speed advantage for small datasets—which however run fast anyway. For larger datasets, this advatage vanishes, and gappa becomes ever more efficient.

### 2 Code Quality

Code quality is an important property of scientific software, as ‘good’ code quality facilitates detecting bugs and ensures long-term maintainability of the software; see [11] for a recent analysis and discussion on the state of software for evolutionary biology. We used the prototype implementation of softwipe (https://github.com/adrianzap/softwipe; also developed in our lab) to assess the relative code quality of genesis. softwipe is a meta-tool for `C` and `C++` software that employs other tools such as clang-tidy and cppcheck to obtain scores for a number of different code quality characteristics/indicators: It checks for compiler warnings, memory leaks, undefined behaviour, usage of assertions, cyclomatic complexity (modularity of the code), code duplication, etc.

The result of softwipe is a ranking of the tested codes from best to worst, scoring each code relative to the others for each code quality indicator, as well as an average “overall” ranking (average over all code quality indicators). At the time of writing this paper, 15 different software codes in `C` and/or `C++` formed part of softwipe code quality benchmark, including highly cited tools such as INDELible [12], MAFFT [13], MrBayes [14], T-coffee [15], and Seq-Gen [16].

Overall, genesis currently achieves the highest score of all tested codes, with 9.6/10 points, with top scores (10/10) in compiler and sanitizer warnings, usage of assertions, and (low) cyclomatic complexity. See https://github.com/adrianzap/softwipe/wiki for details and the up-to-date benchmark results.

**Table S1:** Evaluated libraries and tools for the runtime and memory tests. The table lists the respective programming language, the version we used in our evaluation, the main reference(s), and the accumulated number of citations, using Google Scholar (https://scholar.google.com), accessed on 2019-05-05.

| Tool | Language | Version | Reference | Citations |
| --- | --- | --- | --- | --- |
| GENESIS | `C++` | 0.22.1 | [this article] | - |
| GAPPa | `C++` | 0.5.1 | [this article] | - |
| APE | `R` | 5.3 | [17] | 7390 |
| BIOPYTHON | `Python` | 1.73 | [18] | 1712 |
| DENDROPY | `Python` | 4.4.0 | [19] | 935 |
| ETE3 | `Python` | 3.1.1 | [20] | 644 |
| GGTREE | `R` | 1.10.5 | [21] | 294 |
| GUPPY | `OCaml` | 1.1.a19 | [1] | 476<sup>1</sup> |
| LIBPLL | `C` | 2019-04-13 | [2, 3] | - |
| SCIKIT-BIO | `Python` | 0.5.5 | [22] | - |

<sup>1</sup> GUPPY is part of the PPLACER suite of programs; the number of citations hence also includes those of PPLACER itself.

**Figure S1:**
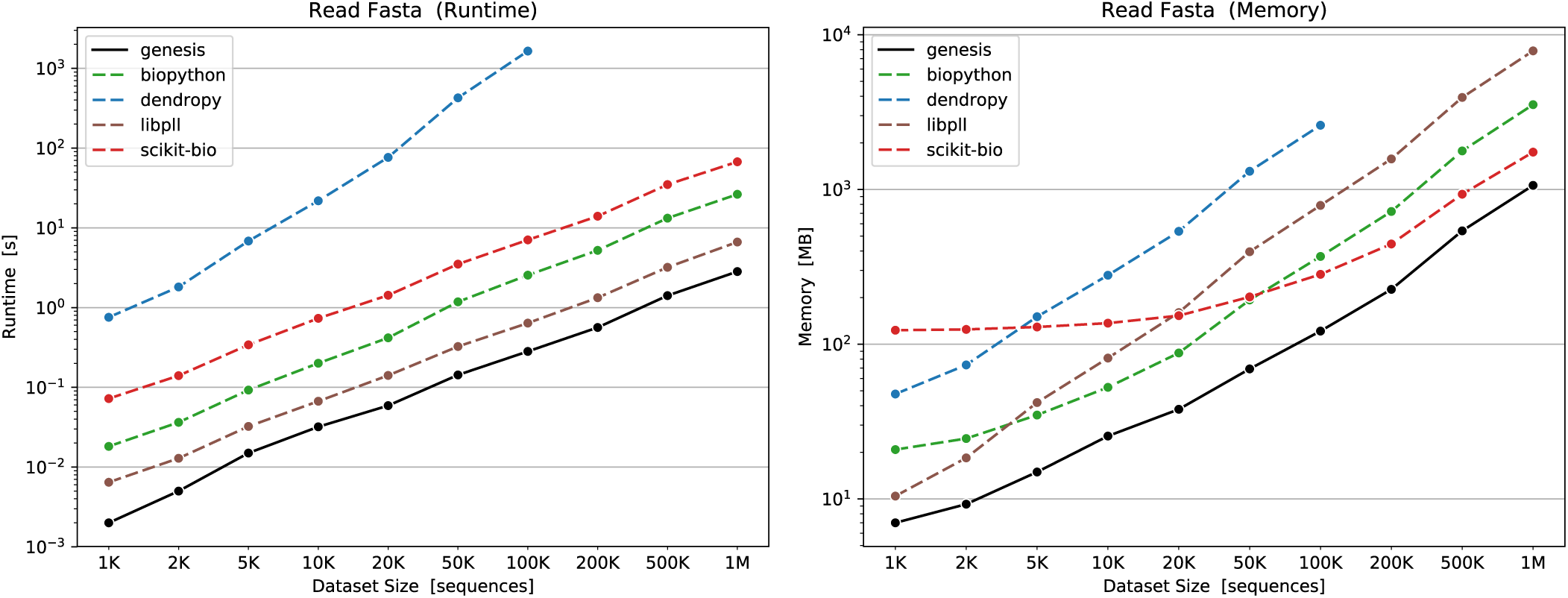
Runtimes and memory footprints for reading fasta files [5] containing 1 K to 1 M sequences. The input files consist of randomly generated sequences using the alphabet ‘`-ACGT`’, with a length of 1000 characters per sequence. Hence, the largest file is about 1 GB in size. In this test, genesis is by far the fastest and most memory-efficient software. In all cases, it parses files more than two times faster than the second fastest software, libpll. Furthermore, the memory usage of libpll for `fasta` files seems to be problematic: It uses about 8 times as much memory as the file size, for example, 7.8 GB for the 1 GB file. On the other hand, genesis has almost no overhead for keeping the files in memory, as can be seen by the 10^3^ MB ≈ 1 GB memory usage for 1 M sequences. Asymptotically, the memory usage of scikit-bio is the second best after genesis: It starts at around 120 MB, probably due to constant memory for its `Python` environment, but then levels out for larger files. However, scikit-bio is also one of the slowest tools, and at least 20 times slower than genesis in all cases. Note that we did not evaluate DendroPy for files larger than 100 K sequences (corresponding to a 100 MB file), because the runtime of 30 min for that file is already impractical for reasonable usage.

**Figure S2:**
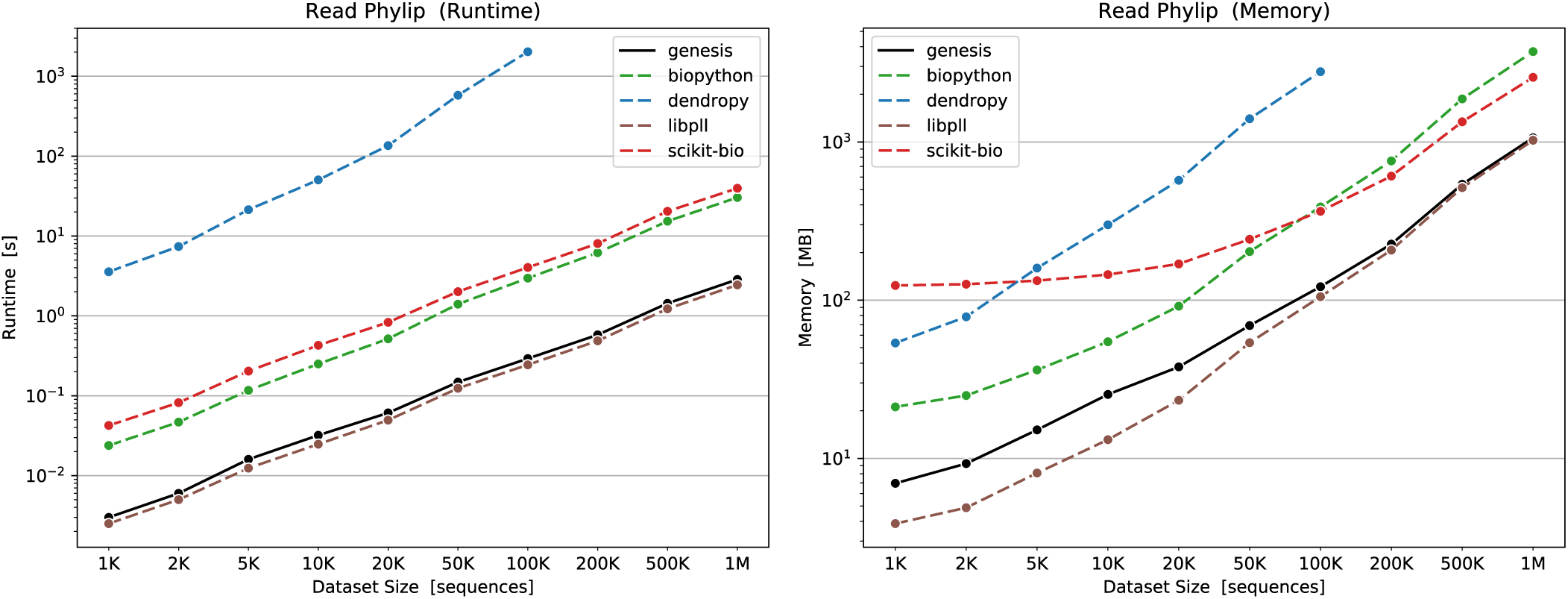
Runtimes and memory footprints for reading phylip files [6] containing 1 K to 1 M sequences. The sequences are the same as in Figure S1, but converted to a strict `phylip` format. Strict `phylip` requires exactly 10 characters for sequence names, and does not allow line breaks in the sequences. This was required by some of the tools, which can not handle “relaxed” `phylip` containing longer sequence names and/or line breaks. Although genesis supports all `phylip` variants, we generally advise against using the `phylip` format for maximum portability across tools. In this test, genesis is second best after libpll in both runtime and memory usage, but only by a small margin. Using the `phylip` format, libpll also does not exhibit the memory issue that it shows for `fasta`, c. f. Figure S1. We again limited the test of DendroPy to reasonable runtimes, as explained in Figure S1.

**Figure S3:**
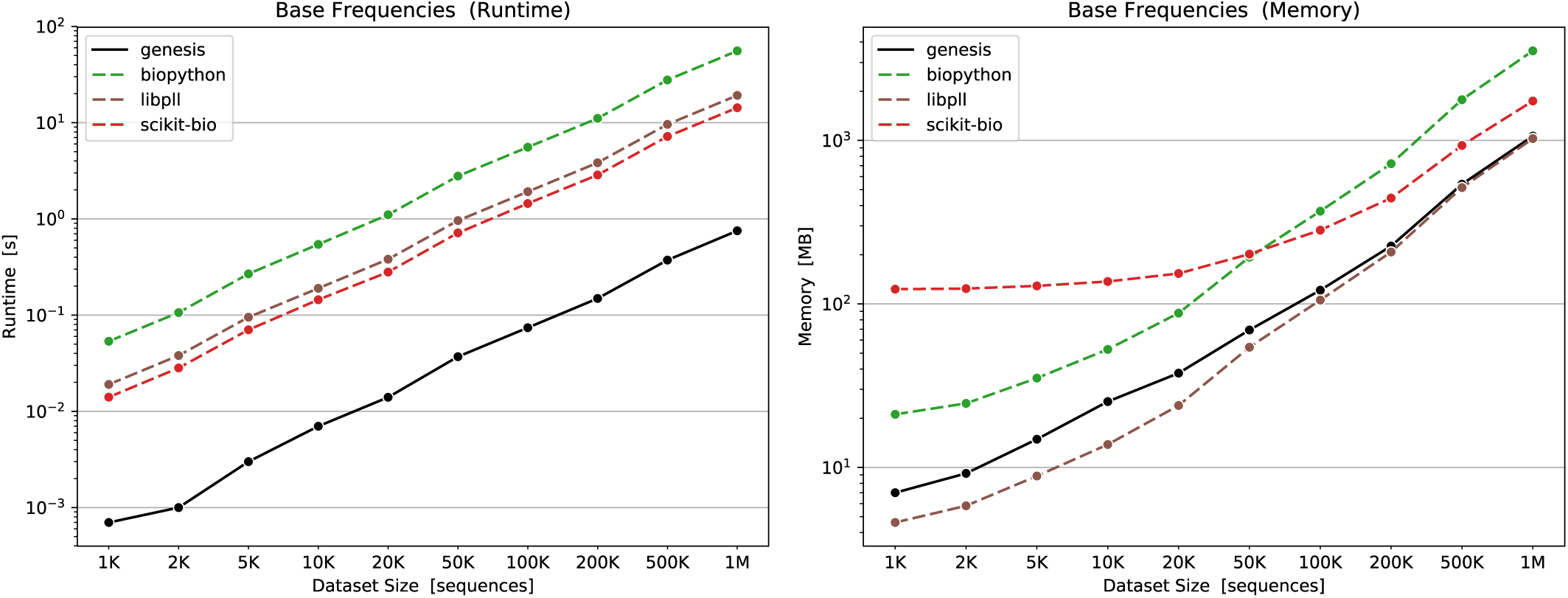
Runtimes and memory footprints for calculating base frequencies on a given set of sequences. Apart from merely reading and parsing files, we also tested some fundamental functionality for working with sequence data. Unfortunately, the range of functions offered by each software largely differs, and we did not find any function or procedure that is provided by all of them. We therefore evaluate the simple calculation of character frequencies/occurences in the sequences of Figure S1 and Figure S2. The parsing of the sequences is *not* included in the time measurement. We only include results for those libraries that do offer a function for counting characters in either single sequences or in an entire set of sequences.

**Figure S4:**
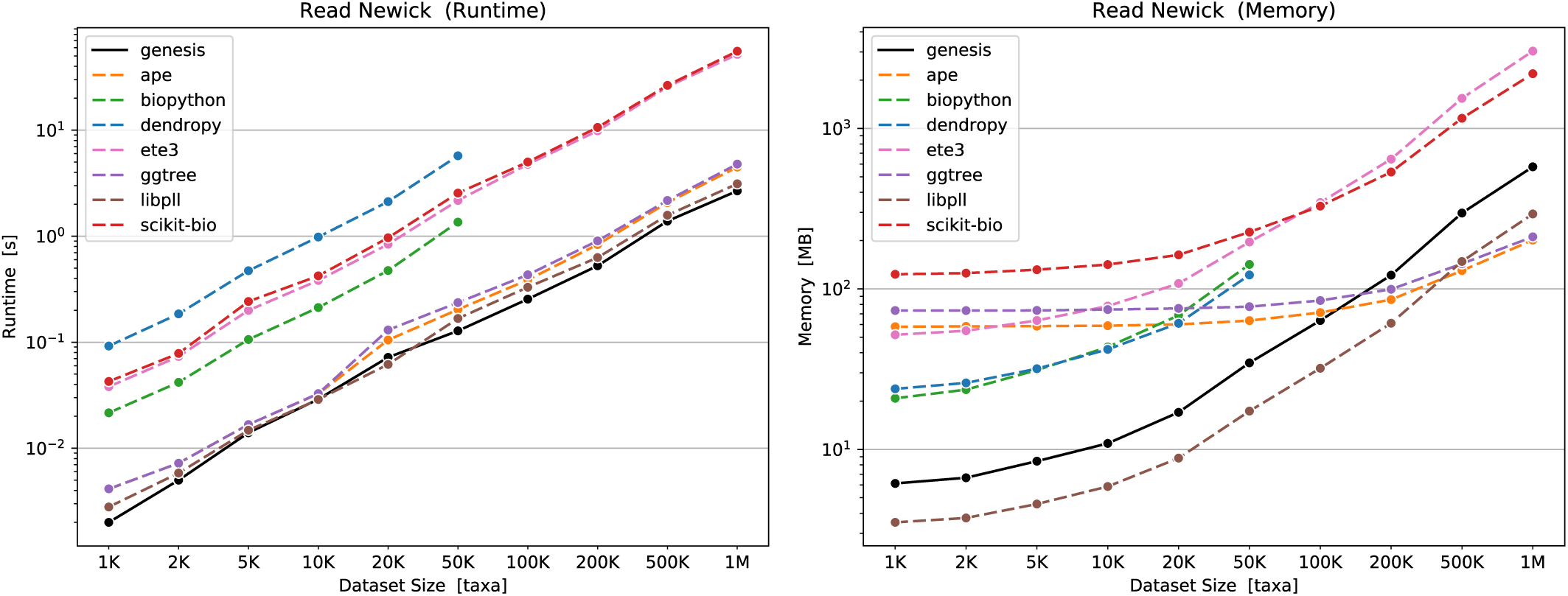
Runtimes and memory footprints for reading newick files [7] with 1 K to 1 M taxa (tip/leaf nodes) and a randomly generated topology. The `newick` format is the only one supported by all libraries that we tested. In most cases, genesis is fastest, closely followed by libpll, which does however exhibit a more efficient memory usage. The memory usage for large trees is best for the `R`-based libraries ape and ggtree, as they store most of the tree data in a memory-efficient table (instead of per-node storage that most other tools use). ete3 exhibits the worst memory efficiency, with 3 GB for a 25 MB `newick` file containing a tree with 1 M taxa. We did not include measurements of DendroPy and Biopython for trees above 50 K taxa. These tools were able to *read* these trees, but not *work* with them: They store and traverse trees recursively, so that even a simple counting of the number of taxa in the tree exceeded the `Python` recursion depth, essentially rendering the tools inapplicable to such large trees.

**Figure S5:**
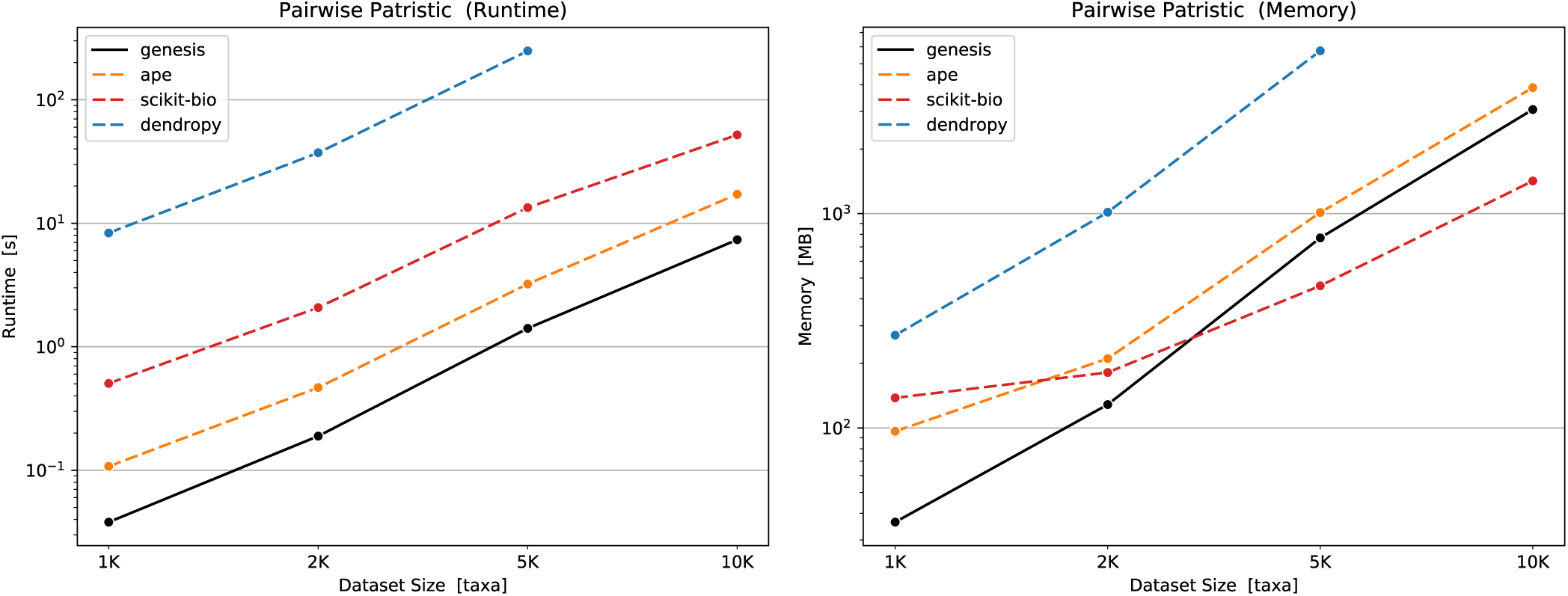
Runtimes and memory footprints for calculating the pairwise patristic distances on a given tree. Similar to Figure S3, we evaluate a simple function that operates on trees: The calculation of the pairwise patristic (branch length) distance matrix between all taxa of the tree. Unfortunately, we could again not identify a basic function that is available in all libraries. For example, ete3 does have a function for calculating distances between two given taxa—but applying this to all pairs of taxa to obtain the full matrix was prohibitively slow, so we did not include it here. The tree parsing times are *not* included in the time measurements. Again, genesis outperforms all other libraries. We note that genesis calculates the matrix for all nodes of the tree, including inner nodes, while the other libraries shown only return the distance matrix for the taxa (leaf nodes) of the tree, which is only a quarter of the size. This explains the better memory footprint of scikit-bio for larger trees. We could not test all tree sizes here, as the resulting matrices would have exceeded the available memory. This is also the reason why DendroPy was not able to calculate the matrix for 10 K taxa on the given hardware.

**Figure S6:**
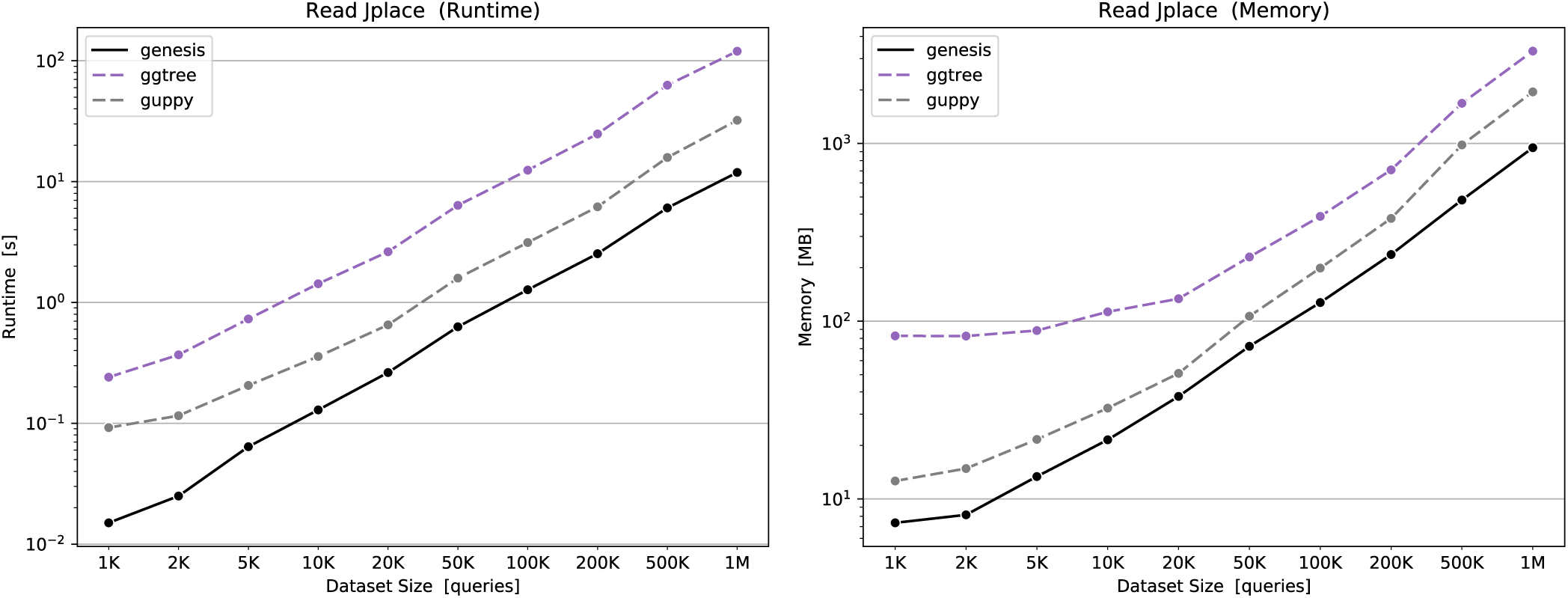
Runtimes and memory footprints for reading jplace files with 1 K to 1 M placed sequences (“queries”). The `jplace` format [8] is a standard file format for storing phylogenetic placements of sequences on a given, fixed reference tree. We conducted this test, because the manipulation, analysis, and visualization of phylogenetic placement data constitutes one of the main use cases of genesis and gappa. Here, we used the 1 K taxon tree of Figure S4 and randomly placed sequences on its branches, with each sequence having up to 5 random placement positions (branches). We include a comparison with guppy, which is a command line tool for the analysis of phylogenetic placements, and the tool on which many ideas and concepts of our gappa tool are based upon. As guppy is not a library, we conduct the test by measuring the runtime and memory usage of its `info` command. The command simply reads a `jplace` file and prints the number of queries stored in the file. genesis consistently outperforms both alternative tools that can parse `jplace` files, and is about one order of magnitude faster than ggtree.

**Figure S7:**
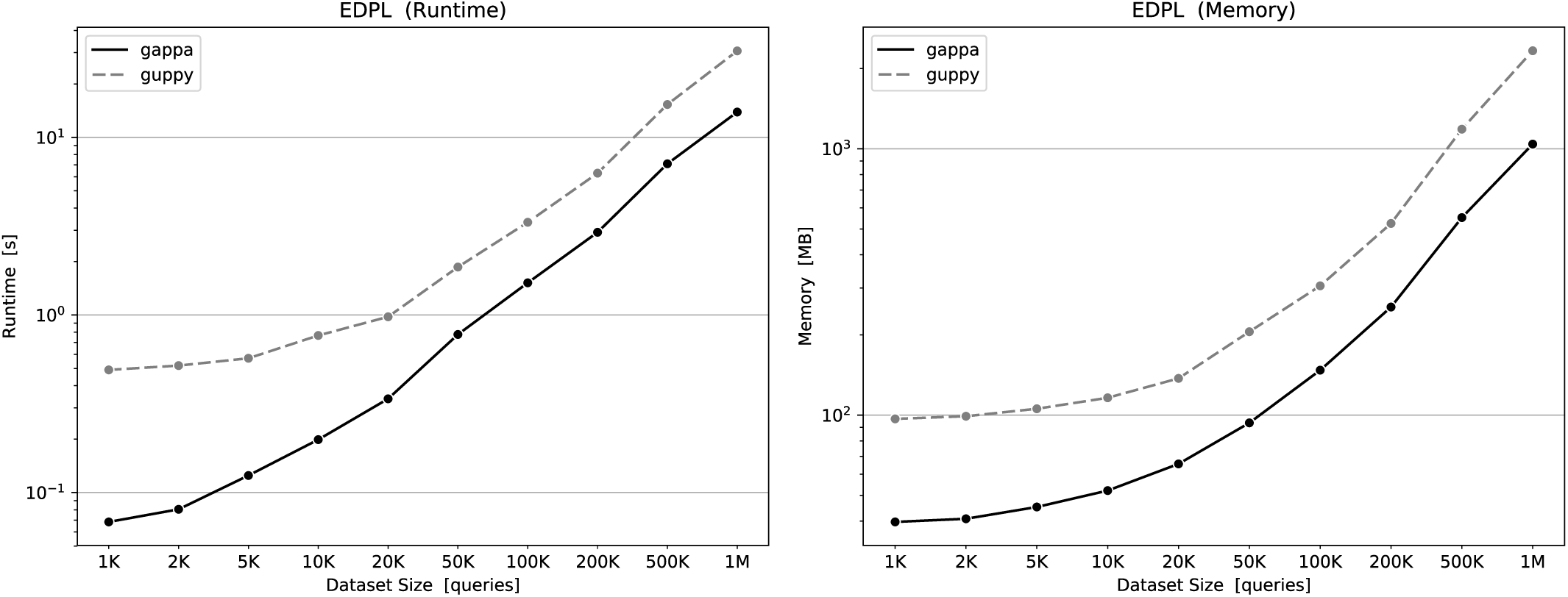
Runtimes and memory footprints for calculating the Expected Distance between Placement Locations (EDPL) [1] of the queries in a jplace file. We again use the randomly generated `jplace` files (so-called “samples”) of Figure S6 with 1 K to 1 M placed sequences (“queries”) here. The EDPL is a metric that measures the uncertainty of each query in a sample, by assessing the distribution of the different placement positions of each query along the branches of the tree. The most time- and memory-consuming part of this analysis is the file reading itself, while the overhead for the actual computation is fairly small. Hence, the plot exhibits similar runtimes and memory footprints as Figure S6. For larger samples, both the runtime and the memory consumption of gappa are about 2 times better compared to guppy.

**Figure S8:**
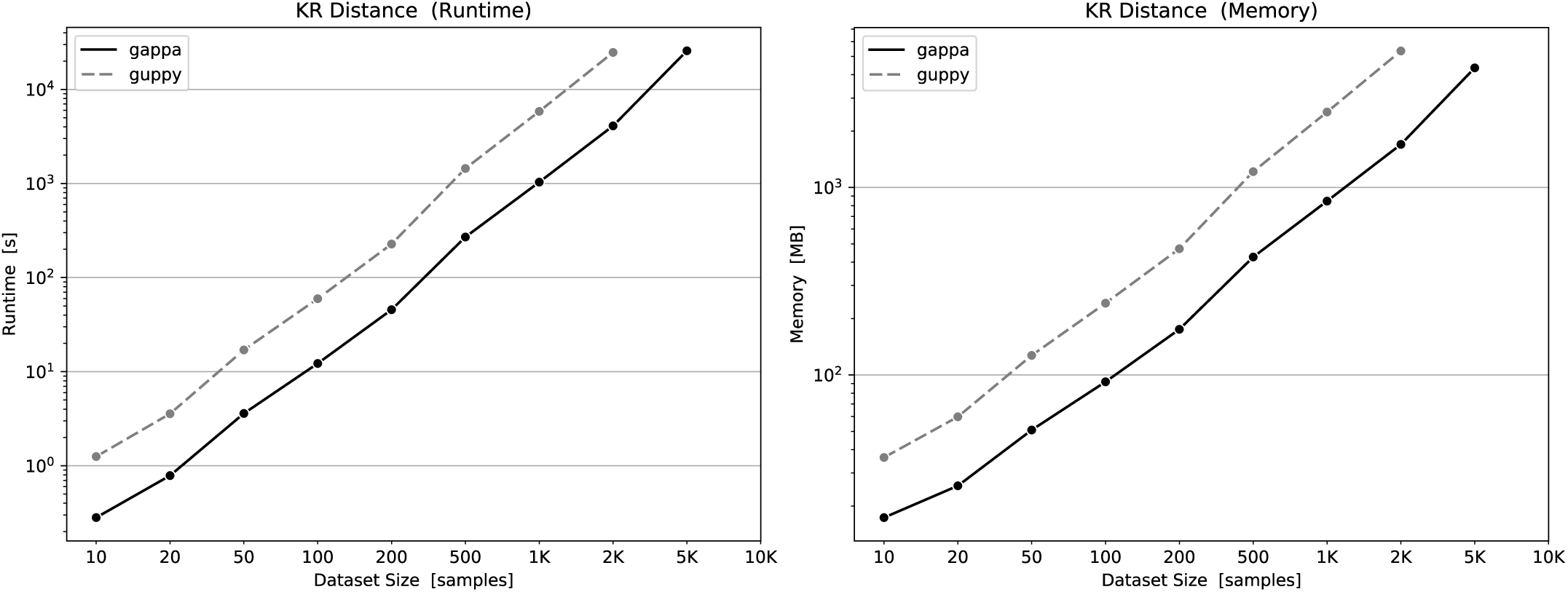
Runtimes and memory footprints for calculating the pairwise Kantorovich-Rubinstein (KR) distance [9] between sets of placement samples. The KR distance is a pairwise metric between placement samples (i. e., between two `jplace` files); hence, we here scale the analysis with the number of samples (`jplace` files). Each sample contains 1000 randomly placed sequences (queries), with each having up to 5 random placement positions (branches). We again used the 1 K taxon tree of Figure S4. The largest analysis with 10K samples corresponds to ≈ 3 GB of `jplace` files; we however could not compute some of the large analyses, as they would have exceeded the available main memory needed for the input data and the resulting pairwise distance matrix. Here, gappa is about 5 times faster, and 3 times more memory efficient than guppy.

**Figure S9:**
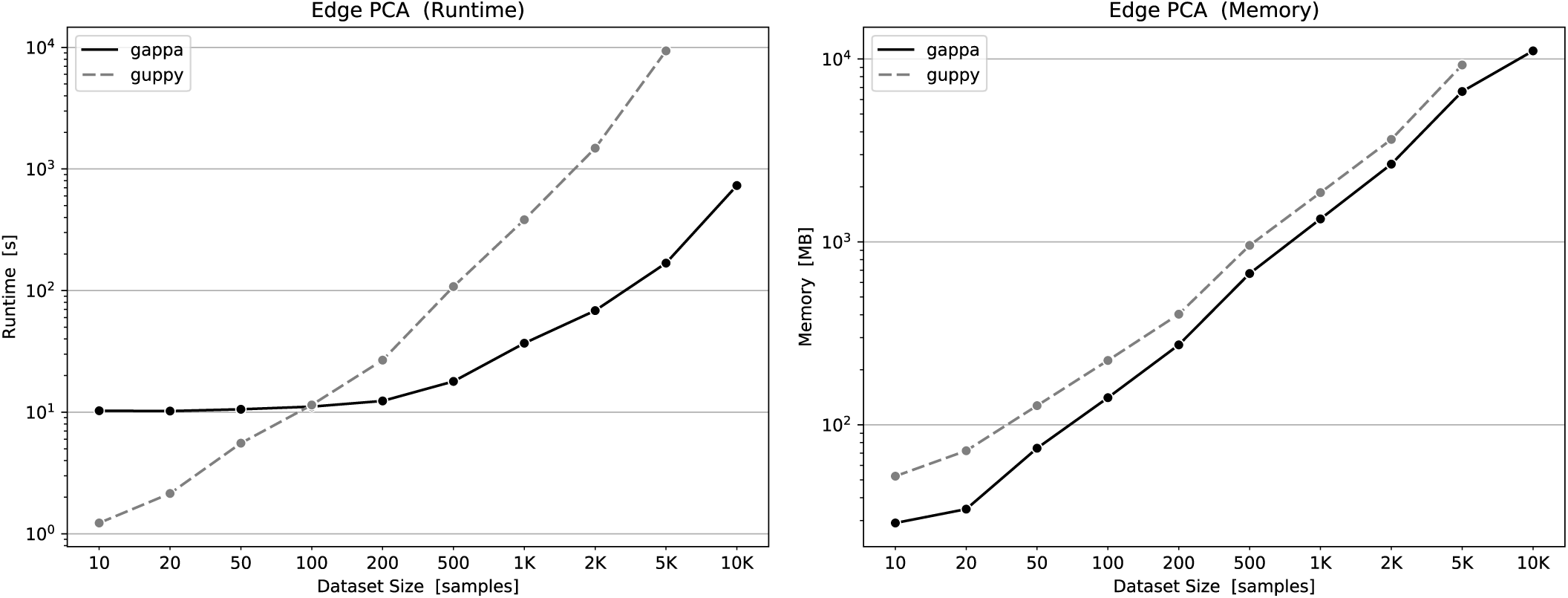
Runtimes and memory footprints for calculating the Edge Principal Components (Edge PCA) of sets of placement samples. The Edge PCA is a typical analysis method for placement data that identifies which taxa of the reference tree are mostly responsible for differences between the composition of distinct samples. Hence, we here again scale the test with the number of samples, again using the random data from Figure S8. The PCA implementation in gappa differs from the one in guppy in that it scales worse with tree size. As the tree size is constant in the test data (1K taxa), we here get a “constant” offset of ≈ 10 s for gappa even for a small number of samples. This however is alleviated by the generally better runtime of gappa that pays off for larger numbers of samples. As the reference trees used in phylogenetic placement usually are around a few thousand taxa is size, this downside of gappa should rarely be an issue in practice. gappa uses about 70% the memory commpared to guppy, and yields growing runtime advantages for larger numbers of samples; for 5K samples, it is 55 times as fast. We could not test guppy with 10K samples due to the limited main memory of 12 GB.

**Figure S10:**
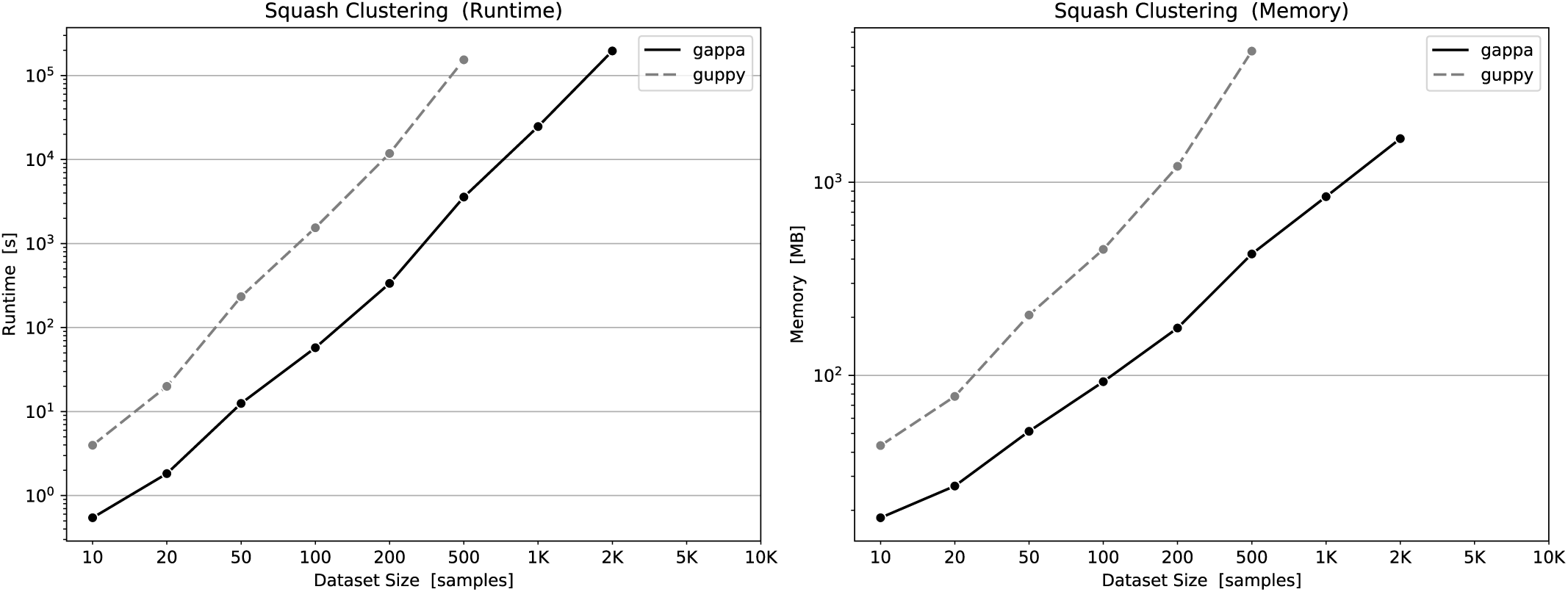
Runtimes and memory footprints for calculating Squash Clustering [10] of sets of placement samples. Squash Clustering is another typical analysis method for placement data that yields a cluster tree of samples by grouping samples that are similar to each other based on their KR distance. We again scale the test with the number of samples, and use the random data from Figure S8. This is by far the most time-demanding type of analysis tested here. The largest tests (with 500 and 2K samples, respectively) correspond to runtimes of ≈ 2 days; we did not run tests with more samples due to these long runtimes. As mentioned, we here limited gappa to run on 1 core only; using more cores yields almost linear speedups. Here, gappa gains larger improvements for larger numbers of samples compared to guppy; for 500 samples, it is 43 times as fast (on a single core) and 11 times more memory efficient.

